# Gradient-based implementation of linear model outperforms deep learning models

**DOI:** 10.1101/2023.07.29.551062

**Authors:** Hamid Hamidi, Dinghao Wang, Quan Long, Matthew Greenberg

**Affiliations:** Department of Mathematics and Statistics; Department of Biochemistry and Molecular Biology; Department of Medical Genetics; Alberta Children’s Hospital Research Institute; Hotchkiss Brain Institute University of Calgary

## Abstract

Deep learning has been widely considered more effective than traditional statistical models in modeling biological complex data such as single-cell omics. Here we show the devil is hidden in details: by adapting a modern gradient solver to a traditional linear mixed model, we showed that conventional models can outperform deep models in terms of both speed and accuracy. This work reveals the potential of re-implementing traditional models with modern solvers.

## Main

Deep learning has made advancements in solving complicated problems and has shown to be effective in discovering complex nonlinear structure in high-dimensional data^1^.

Deep learning models require large amounts of data in order to learn such structure. Working with large datasets requires scalable algorithms. Powerful, open source software packages like PyTorch^2^ and Tensorflow^3^ provide flexible languages for constructing deep learning models as well as implementations of scalable algorithms for fitting them, making deep learning techniques extremely accessible to researchers. This accessibility, together with the high-profile success of deep learning in computer vision and natural language processing, has led researchers to adopt deep learning for biological data analysis.

Although deep learning has become popular for modeling single-cell gene expression data, its performance relative to traditional statistical models has not been fairly investigated as people use modern solvers for deep learning and traditional solvers for traditional models. In this work, we compare the performance of scVI, a deep learning model^4^, with that of ZINB-WaVE, a traditional statistical model^5^, both using modern solvers. By virtue of the powerful software used to implement it, scVI scales well to large datasets. On the other hand, the bespoke algorithm designed to fit the ZINB-WaVE model has inferior scaling properties and cannot be applied to large datasets. To effect a rigorous comparison of scVI and ZINB-WaVE in the setting of large data, we develop a scalable algorithm, reminiscent of alternating least squares, for fitting ZINB-WaVE models. In implementing this algorithm, we borrow the stochastic gradient descent-based model fitting machinery used in deep learning. Our subsequent analysis facilitated by this implementation suggests that ZINB-WaVE matches the performance of scVI both on goodness-of-fit and in performance on downstream tasks even though it is comparatively simple. Moreover, being linear, the ZINB-WaVE model is interpretable whereas, like most deep learning models, scVI is a “black box”. Interpretability is highly desirable in the biological and especially in medical contexts. For these reasons, we recommend ZINB-WaVE, rather than scVI, for modeling single-cell gene expression data.

Current experimental limitations such as high variance, sparsity, limited and variable sensitivity, batch effects, and transcriptional noise^6,7^ give rise to unique challenges in scRNA-seq data. In recent years, various computational tools have been developed to overcome these challenges^6^. Models such as DCA^8^, scVI^4^, ZINB-WaVE^5^, and scDCC^9^, all based on Zero-Inflated Negative Binomial (ZINB) distribution, have shown superior performance compared to models based on other distributions (e.g., MAGIC^10^, BISCUIT^11^, and ZIFA^12^). In this work, we focus on ZINB-WaVE^5^. Introduced in 2017 and, since, ascendant among the ZINB-based models, ZINB-WaVE is a generalized linear mixed model that combines dimensionality reduction, batch effect correction, and technical variability (e.g. dropouts) all in one model. It captures technical variability through two random variables and maps the data onto a biologically meaningful low-dimensional representation through a linear transformation^5^. One of the descendants of ZINB-WaVE^5^ is the deep learning-based scVI^4^ model, published in late 2018. It uses a variational autoencoder whose latent variables are specified by normal distributions. It generates estimates for the parameters of the ZINB distribution via a nonlinear transformation of transcriptomes^4^ by a neural network. Lopez *et. al*. showed that scVI captures meaningful biological information and scales to a million cells^4^.

By contrast, the iterative algorithm developed in 2018 for fitting ZINB-WaVE models cannot be scaled to more than 30,000 cells^4^. This is unsurprising since this algorithm repeatedly applies ridge and logistic regression, SVD on matrices with dimension equal to the sample size, and the BFGS quasi-Newton method^2^. To overcome this limitation, we developed a stochastic gradient descent-based optimization procedure for fitting ZINB-WaVE^5^ models and coded it in PyTorch yielding a scalable, high-performance, GPU accelerated, memory-efficient algorithm and memory-efficient implementation (**Online Methods**). Armed with this algorithm, we fit the ZINB-WaVE model to various large benchmarking data sets, including CORTEX^13^, RETINA^14^, and BRAIN^15^, We assessed both the performance of our model fitting algorithm and the quality of the resulting fit models (**Supplementary Table S1**).

To quantify ZINB-WaVE’s scalability and efficiency, we used randomly selected cells from the BRAIN^15^ data set with 720 highly-variable genes across all cells (**Online Methods**). Our analyses showed that ZINB-WaVE’s run-time is significantly lower than that of scVI^4^ for any data size (**Fig. 1A**). This is entirely unsurprising, due to the relative simplicity of the ZINB-WaVE model. We used the loss function, negative log-likelihood, evaluated on the training (resp. testing) set to compare the goodness-of-fit (resp. generalization) of ZINB-WaVE and scVI. On the BRAIN data set, ZINB-WaVE performed better than scVI in terms of goodness-of-fit and generalization. (**Fig. 1B**). For the CORTEX^13^ and RETINA^14^ data sets, the average negative log-likelihood of ZINB-Grad is the same or comparable to scVI^3^ and ZINB-WaVE^2^ (**Fig. 1C** and **Supplementary Fig. S2**).

**Figure 1:**
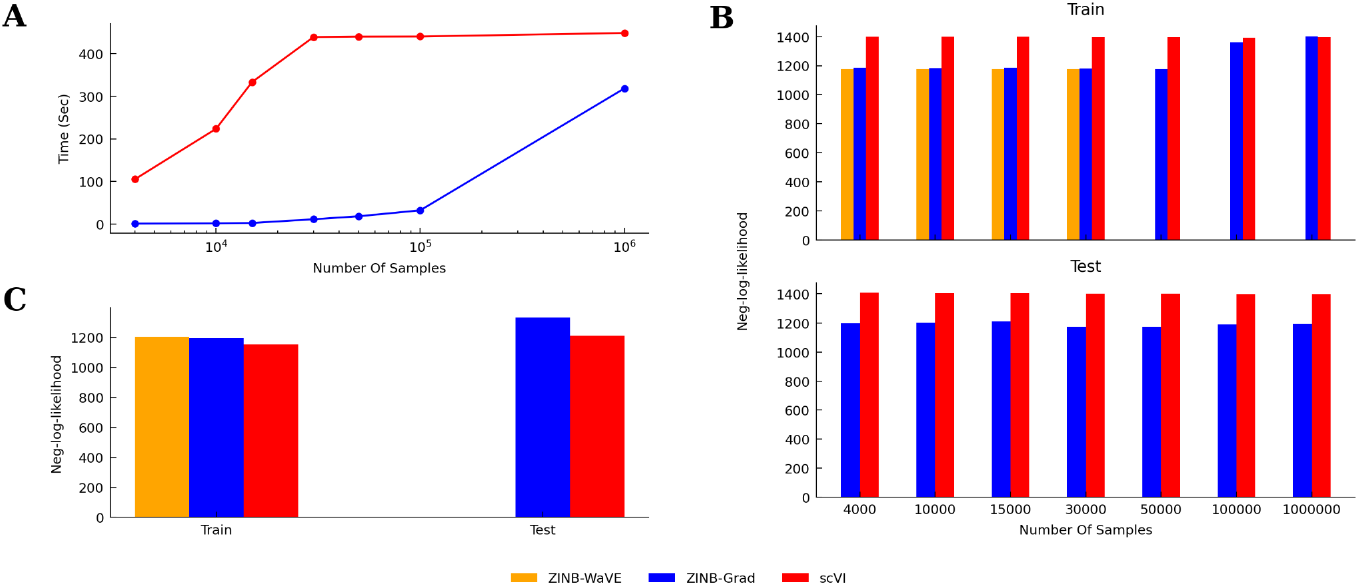
The metrics used and ZINB-Grad Performance. (A) Run-times for inference parameters of the BRAIN data set. (B) Average negative log-likelihood (the smaller the better) on the training data and the test (held-out) data from the BRAIN data. (C) Average negative log-likelihood on the train and test from the CORTEX data set We then assessed the ZINB-Grad latent space by examining biological variation explained in the latent space using CORTEX^13^ and RETINA^14^ data sets and unsupervised cell clustering.

We used three measurements, namely, Normalized Mutual Information (NMI), Adjusted Rand Index (ARI), and Average Silhouette Coefficient (ASC) to compare the performance of clustering (**Online Methods**). The CORTEX^13^ data set clustering scores for scVI^4^ and ZINB-Grad show that ZINB-Grad performed slightly better than scVI^4^ (**Fig. 2A**). ZINB-Grad and ZINB-WaVE^5^ accuracy measurements are close for both CORTEX^13^ and RETINA^14^ data sets (**Fig. 2A** and **Supplementary Fig. S1**).

**Figure 2:**
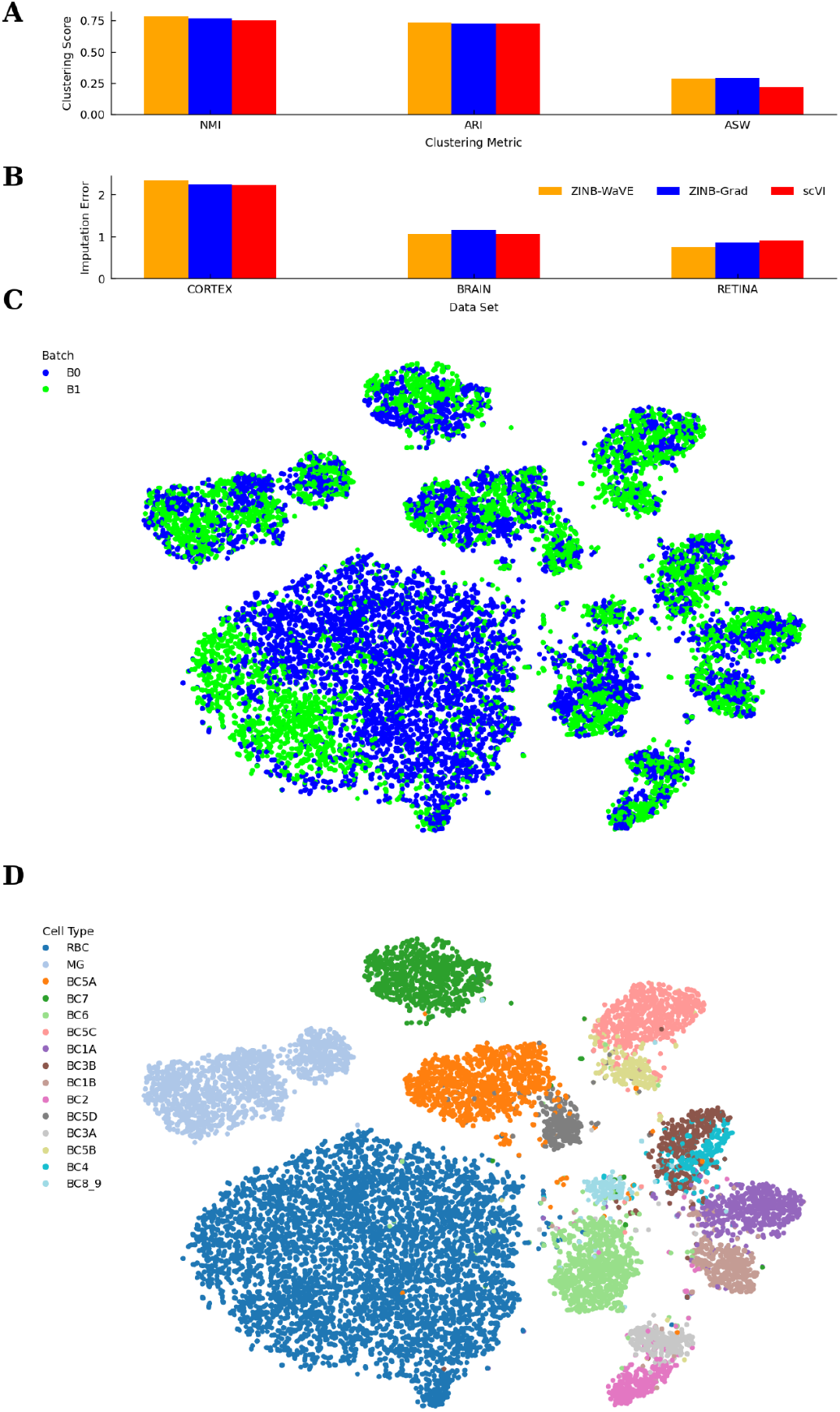
The ZINB-Grad Performance (continued). **(A)** Clustering accuracy for the CORTEX data set using NMI, ARI, and ASW. **(B)** Imputation error for the CORTEX, RETINA, and BRAIN data sets. **(C)** The visualization of the latent space of ZINB-Grad for the RETINA data set after batch correction. **(D)** Same latent space with cell types.

We further assessed ZINB-Grad using all three data sets^13–15^ (**Fig. 2B**) on imputation error between original and imputed data (**Online Method**). For all data sets, the ZINB-Grad imputation error is better or comparable to scVI^4^ and ZINB-WaVE^5^ (**Fig. 2B**). We evaluated the accountability for technical variability by assessing batch entropy of mixing (**Online Methods**) and visualizing the latent space (**Fig. 2C,D**) in the RETINA^14^ data set containing two batches. ZINB-Grad entropy of batch mixing is better than ZINB-WaVE^5^ and comparable to scVI^4^ performance (**Supplementary Fig. S3**). These results show that ZINB-Grad has biologically meaningful latent space performing as good as scVI^4^ and ZINB-WaVE regarding data imputation and accountability for technical variations.

Sc-RNA-seq technology has revolutionized scientists’ viewpoints on many complex biological processes such as embryo development and tumor microenvironment. With an incremental trend in the amount and scale of single-cell experiments, the need for an efficient and accurate tool in analyses of sc-RNA-seq data is becoming vital. There is a tendency toward developing deep models for sc-RNA-seq experiments, which might be used cautiously. Our development shows that a conventional model optimized with the proper techniques and implemented using the right tools can outperform state-of-the-art deep models. It can generalize better for unseen data and uses substantially fewer resources compared to deep models.

## Acknowledgement

QL is partly supported by the NSERC Discovery Grant (RGPIN-2017-04860) and the New Frontiers in Research Fund (NFRFE-2018-00748). DW is supported by the Alberta Graduate Excellence Scholarship.

## Author Contributions

Conceived the study: MG. Developed the tool: HH, MG. Analyzed data: HH, DW, QL, MG. Supervised the study: QL, MG. Wrote the manuscript: HH, QL, MG.

## Online Methods

### The ZINB-Grad model and its gradient-based estimation

**The ZINB-Grad model** adapts the ZINB-WaVE^5^ model with zero-inflated negative binomial (ZINB) distribution. The probability mass function (PMF) of the negative binomial distribution with mean *µ* and inverse dispersion parameter *θ* is:

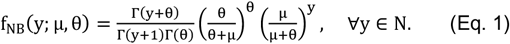

With π ∈ [0,1], which represents the probability that a 0 is observed instead of the actual count, and δ_0_(.) as the Dirac function, the ZINB distribution PMF is as follows:

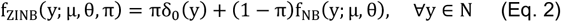

The π in (Eq. 2) is essentially the probability of dropouts which is a common issue in sc-RNA-seq data. Therefore, the ZINB distribution can be effective in modeling the dropouts.

Given *Y* as the matrix of observations with *n* samples and *j* features, Y_ij_ denotes the count of feature j in sample i. Risso *et. al*. modeled the Y_ij_ as a random variable following ZINB distribution. They considered *µ* _ij_, *θ* _ij_ and *π*_*ij*_ as parameters with the following linear mixed effect model:

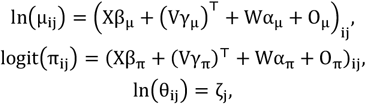

Where logit 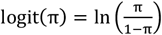

*X* and *V* are known sample-level and gene-level covariates, respectively. *X*can model wanted or unwanted variations. For instance, *X*can be used to correct batch effects, and *V* can model gene length or GC content.

*W* and α are the unknown matrices that are used for the dimension reduction part. *W* is essentially the latent space of ZINB-WaVE, and α is its corresponding matrix of regression parameters. The *O* parameters are the known offsets for *π* and *µ* . For more details, please refer to ZINB-WaVE^2^.

Here, we considered the same *W* and *α* but different *V* and *X*for *π* and *µ* .

### The estimation procedure of ZINB-Grad

The parameters to be inferred are *β, γ, W, α*, and *ζ*, and known parameters are *X, V, Oπ*, and *Oπ*. Given the data (count matrix *Y*), the log-likelihood function is:

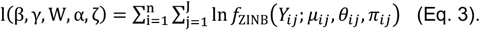

We use a regularized maximum log-likelihood estimation (MLE) approach to infer the parameters. However, instead of maximizing the log-likelihood, we minimize the negative log-likelihood and estimate the parameters by trying to solve:

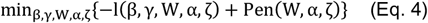

where pen(.) is the regularization term to prevent overfitting as follows:

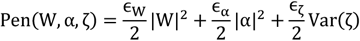

| ⋅| is the Frobenius matrix norm and (*ϵ*_W_, *ϵα, ϵα*) are non-negative regularization parameters.

We randomly initialized the parameters and set (Eq. 4) as the objective function. We used gradient descent, backpropagation, and *Adam* optimizer (a first-order stochastic optimizer^16^) with a learning rate of 0.08 to minimize the complex non-convex objective function. We optimized the objective function until convergence -usually 150 iterations, where each iteration is the complete pass through the data set.

For larger data sets (with >15,000 samples), we used a two-step optimization procedure for memory and performance efficiency as follows:

Step1 - a complete pass through the whole data set with batches of a maximum size of 10,000 samples. In this step, the inference procedure will estimate a global *β, α*, and *ζ* for all samples. Since our main goal in this step is to find the best *β, α*, and *ζ* for all samples, we used an adaptive number of iterations to have fixed optimization steps for these parameters and avoid overfitting. We used these parameters along with *W* and *γ* corresponding to each sample for the next step.

Step2 - Another complete pass through the whole data set with the same batches from the first step. In this step, since we already found the best global *β, α*, and *ζ* in the previous step, we do not optimize them any further (freeze them). However, we optimize for *W* and *γ*, which need fewer iterations (usually 30 iterations) since they have been optimized in the first step.

### Datasets and comparisons

#### Data sources

CORTEX consists of 3,005 mouse cortex cells and gold-standard labels for seven different cell types from reference^13^. As described by their original publications refs. ^4,11^, we used 558 high-variable genes as the features.

RETINA is the data set from ref.^15^ with filtering process and cluster annotations from the original study.

BRAIN is a dataset of 1.3 million brain cells from 10x Genomics^15^. We selected 720 highly-variable genes across all cells, as described in ref.^4^. We randomly shuffled the data to get a subset of the samples for efficiency and generalization analyses.

#### Run-time and generalization

To evaluate the efficiency and goodness of fit, we randomly subsampled the BRAIN data set with different data sizes (of 4K, 10K, 15K, 30K, 50K, 100K, and 1M). We used run-time and negative log-likelihood to estimate efficiency and the goodness-of-fit, respectively. For this purpose, we used the negative log-likelihood of the test data (held-out data or unseen data) in addition to the training data. The validation of the ZINB-Grad is performed by fixing *β, α*, and *ζ* learned from the training set and inferring *W* and *γ* through a few iterations (default is 20). All scVI and ZINB-Grad tests were performed using a computer equipped with a V100 GPU and 32 GB of system memory.

#### Imputation Evaluation

We adopted the imputation benchmarking introduced in ref.^4^. We randomly selected 10% of the non-zero entries and altered them to zero with a Bernoulli process of probability 0.9. Then, we used the median of the L_1_ distance between the original and imputed values of the altered data set as the accuracy for data imputation.

#### Clustering

We performed clustering on the latent space with 10 dimensions for scVI^4^, ZINB-WaVE^5^, and ZINB-Grad using K-means with 50 different random initialization and 400 iterations. We used three different clustering metrics, namely, Normalized Mutual Information (NMI), Adjusted Rand Index (ARI), and Average Silhouette Coefficient (ASC) to evaluate the performance.

Assume U and P as the categorical distributions of real labels and predicted labels, respectively. The NMI is as follows:

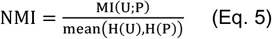

where H(.) is the Shanon entropy and *MI* is the mutual information.

We calculated the NMI (Eq. 5) between the gold standard labels and labels obtained from K-means to assess the clustering performance. The NMI is between 1.0 and 0. The former shows perfect clustering, and the latter shows no mutual information.

The ARI is defined as follows:

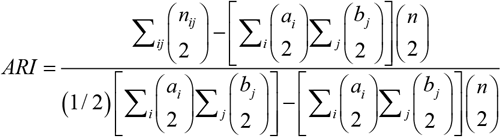

where *n*_*ij*_, *a*_*i*_, *b*_*/*_ are values from the contingency table.

Furthermore, for a data point i the Silhouette Coefficient is defined as 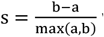 where *a* is the mean distance between sample i and all other points in the same cluster, and *b* is the average distance between sample i and all other points in the next nearest cluster. We considered the average Silhouette Coefficient for all samples as ASC. The perfect ASC is equal to 1.

#### Batch effect correction

We used the entropy of batch mixing to assess the batch effect correction as in refs.^4,17^. We randomly selected 100 cells from batches and found 50 nearest neighbors of each randomly chosen cell to calculate the average regional Shanon entropy of all 100 cells. The procedure is repeated for 100 iterations, and the average of the iterations is considered as the batch mixing score.

## SUPPLEMENTARY MATERIALS

**Supplementary Figure S1:**
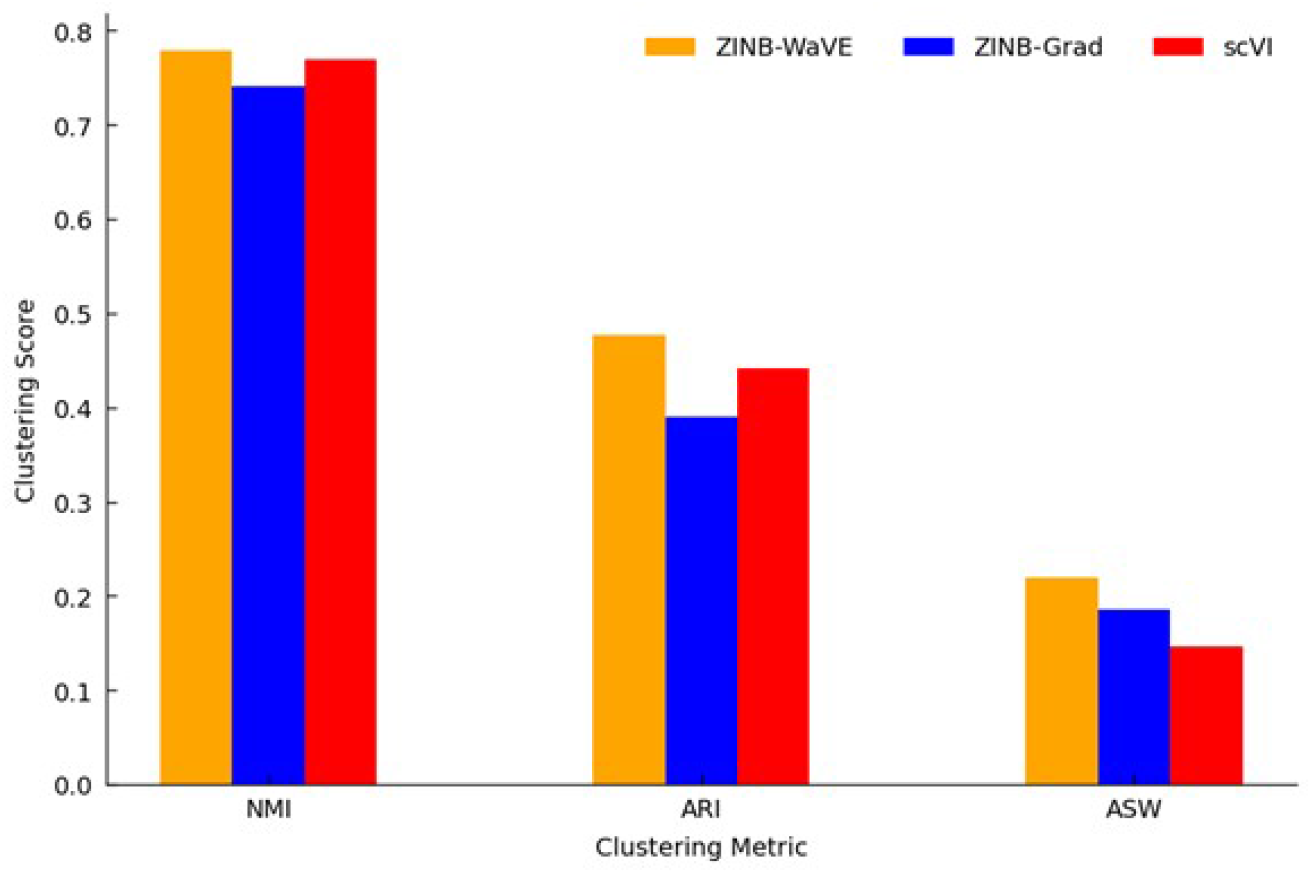
Various clustering metrics for the RETINA data set.

**Supplementary Figure S2:**
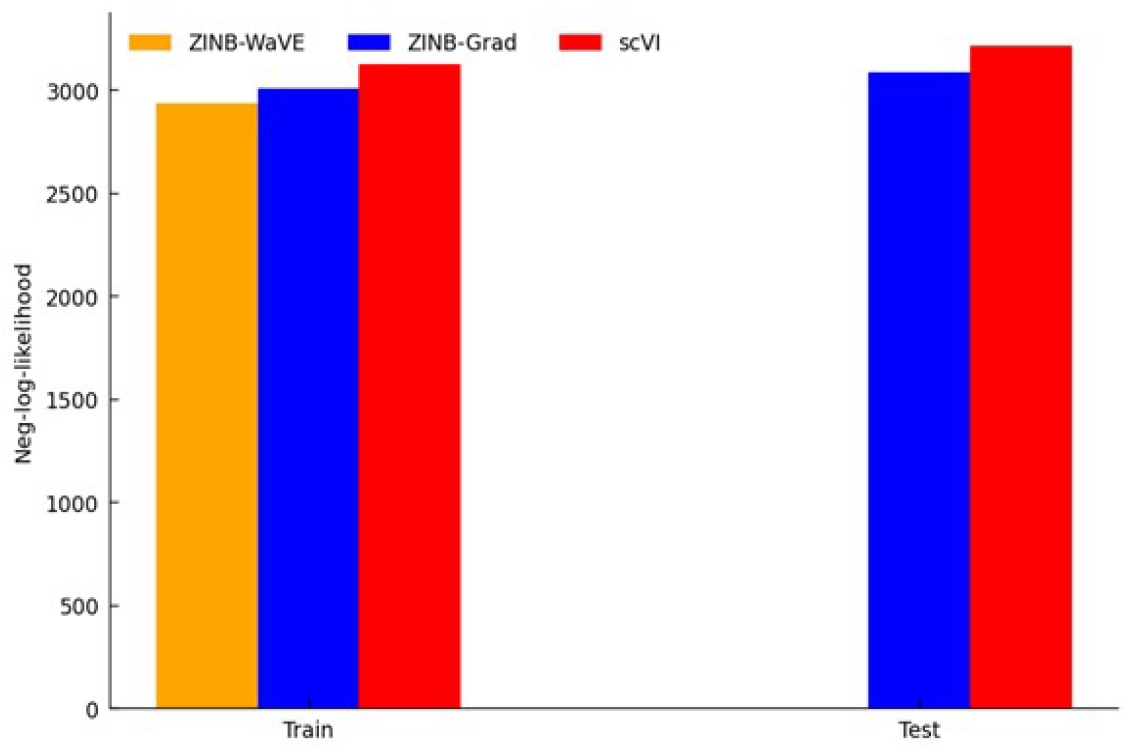
Average negative log-likelihood (the smaller the better) on the training data and the test (held-out) data from the RETINA data set.

**Supplementary Figure S3:**
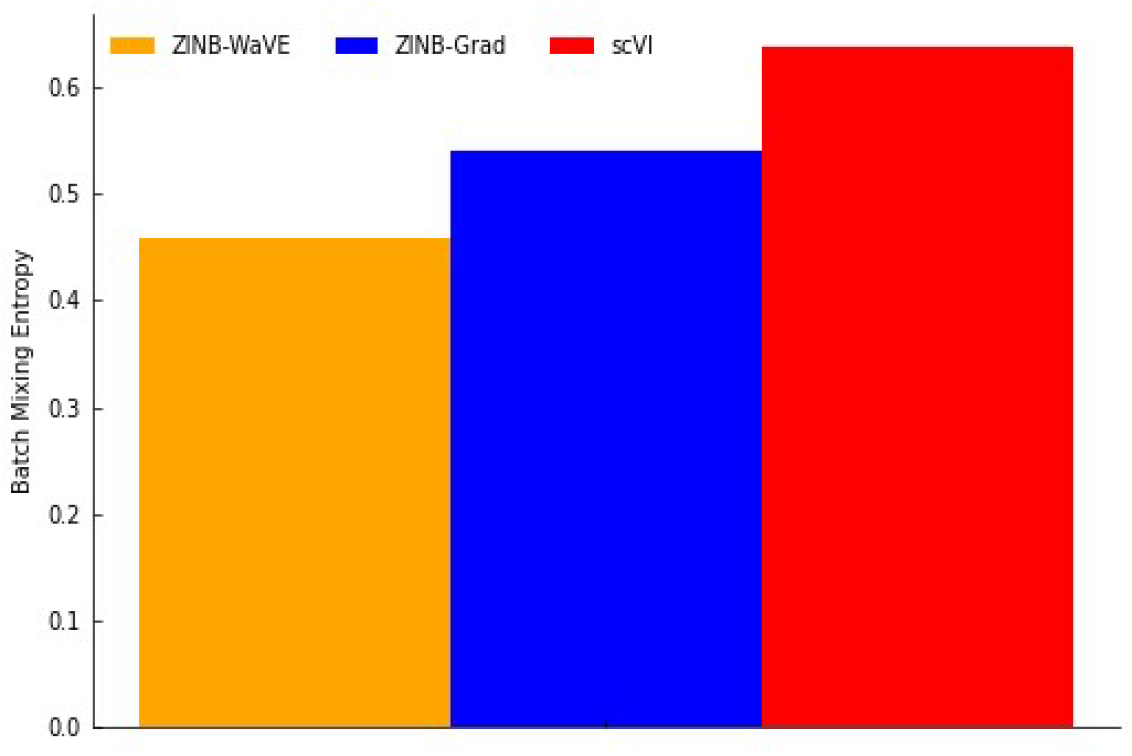
Entropy of Batch Mixing for the RETINA data set (the higher, the better).

**Supplementary Table S1:**
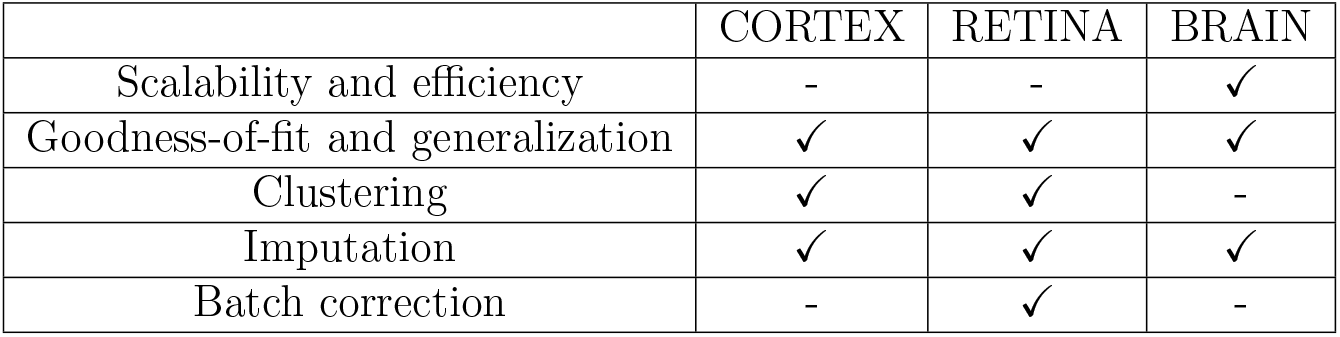
The benchmark tests and data sets used for comparing ZINB-Grad and other models.

|  | CORTEX | RETINA | BRAIN |
| --- | --- | --- | --- |
| Scalability and efficiency | - | - | ✓ |
| Goodness-of-fit and generalization | ✓ | ✓ | ✓ |
| Clustering | ✓ | ✓ | - |
| Imputation | ✓ | ✓ | ✓ |
| Batch correction | - | ✓ | - |

## Notes

### Competing Interest Statement

The authors have declared no competing interest.

